# Cell-penetrating peptide and cationic liposomes mediated siRNA delivery to arrest growth of chronic myeloid leukemia cells *in vitro*

**DOI:** 10.1101/2023.12.27.573407

**Authors:** Vera Vysochinskaya, Yana Zabrodskaya, Olesya Dovbysh, Anton Emelyanov, Vladimir Klimenko, Nikolay Knyazev, Ivan Terterov, Marya Egorova, Alexey Bogdanov, Michael Maslov, Andrey Vasin, Michael Dubina

## Abstract

Gene silencing through RNA interference (RNAi) is a promising therapeutic approach for a wide range of disorders, including cancer. Non-viral gene therapy, using specific siRNAs against *BCR-ABL*, can be a supportive or alternative measure to traditional chronic myeloid leukemia (CML) tyrosine kinase inhibitor (TKIs) therapies, given the prevalence of clinical TKI resistance. The main challenge for such approaches remains the development of the effective delivery system for siRNA tailored to the specific disease model.

The purpose of this study was to examine and compare the efficiency of endosomolytic cell penetrating peptide (CPP) EB1 and PEG_2000_-decorated cationic liposomes composed of polycationic lipid 1,26-bis(cholest-5-en-3-yloxycarbonylamino)-7,11,16,20-tetraazahexacosane tetrahydrochloride (2X3) and helper lipid 1,2-dioleoyl-sn-glycero-3-phosphoethanolamine (DOPE) for anti-bcr-abl siRNA delivery into the K562 human CML cell line. We show that both EB1 and 2X3-DOPE-DSPE-PEG_2000_ (0.62% mol.) liposomes effectively deliver siRNA into K562 cells by endocytic mechanisms, and the use of liposomes leads to more effective inhibition of expression of the targeted gene (*BCR-ABL*) and cancer cell proliferation. Taken together, these findings suggest that PEG-decorated cationic liposomes mediated siRNA delivery allows an effective antisense suppression of certain oncogenes, and represents a promising new class of therapies for CML.

## 1. Introduction

Chronic Myeloid Leukemia (CML) is a type of clonal myeloproliferative hematopoietic stem cell disorder, characterized by the presence of the Philadelphia chromosome (Ph), which results in the formation of the BCR–ABL fusion oncoprotein. This fusion protein exhibits deregulated tyrosine kinase activity [1], which has been shown to be necessary and sufficient for the transformed phenotype of CML cells [2]. The development of small-molecule targeted drugs known as tyrosine kinase inhibitors (TKIs) was designed to interfere with BCR-ABL tyrosine kinase activation by competitive binding at the ATP-binding site [3][3][4]. TKI treatment has resulted in significant breakthrough in the therapy of patients with CML. However, TKI have different potencies, sensitivities to kinase domain mutations, cross-reactivity with other kinases, and patients treated with these newer agents have experienced inadequate responses and early relapses [5]. As a result, there is an urgent need to develop novel therapeutics for CML patients.

The discovery of the RNA interference mechanism fueled the development of small interfering RNA (siRNA)-based drugs for treating various diseases, including cancer [6][7]. Synthetic siRNA can be delivered into target cells to degrade desired mRNA for down-regulate specific proteins involved in the pathogenesis of the disease. The siRNAs provide an alternative approach to reducing the effects of *BCR-ABL* transcripts by regulating oncogene mRNA levels [8]. Moreover, combining TKI with siRNA against *BCR-ABL* may potentiate the anti-tumor effect [9][10]. However, the main challenge in translating BCR-ABL siRNA-based drugs into clinical practice is developing an efficient siRNA delivery system that can specifically target cells.

Cell-penetrating peptides (CPPs) have emerged as promising delivery vehicles and have been demonstrated to be an effective siRNA delivery system in various *in vitro* and *in vivo* tumor models [11][12][13]. CPPs are short, typically positively charged peptides with specific membrane active properties that facilitate their internalization along with covalently or noncovalently complexed cargo-molecules [13]. The mechanism of siRNA delivery by CPP remains unclear and depends on experimental conditions[14]. As a result, the common approach of using CPP/siRNA complexes as universal delivery system may not be suitable for different cell types. In present study, we examined the endosomolytic CPP EB1 [15]as a delivery system for BCR-ABL siRNA. We hypothesized that early endosomes in K-562 cells, which are more acidic than those in other cell types[16], would provide an advantage for effective endosomal escape and target gene silencing.

Another promising component that represents liposomal siRNA delivery vehicles is polycationic amphiphiles based on cholesterol and natural polyamines. The main structural component of such liposome, 1,26-bis(cholest-5-en-3β-yloxycarbonylamino)-7,11,16,20-tetraazahexacosane tetrahydrochloride (2X3), consist of two structural domains: hydrophobic (cholesterol) and hydrophilic polycationic (spermine), which are connected by oligomethylene spacer groups using a carbamate linker. Due to their positive charge, liposomes based on 2X3 can form complexes with negatively charged nucleic acids. These delivery systems have shown their efficacy both *in vitro* and *in vivo* [17] [18][19][20][21].

Currently, lipid-based nanoparticles (LNPs) are primarily used for delivering nucleic acid in clinical practice [22][23]. Although LNPs offer a promising opportunity for efficient intracellular nucleic acid delivery, challenges still remain [24]. The development of new delivery platform technology is crucially required, particularly for systemic administration, which is required for many oncological diseases, including CML.

In present study, we tested and compare two different non-viral cargoes for anti-bcr-abl siRNA delivery and their antiproliferative activity on K-562 cells (CML in myeloid blast crises). The first system was based on CPP EB1, while the second was based on cationic liposomes 2X3-DOPE (1:3 mol.) modified by DSPE-PEG_2000_ (0.62% mol.) (2X3-DOPE-PEG). Our results showed that both siRNA delivery vehicles effectively silenced the *BCR-ABL* gene and inhibited cell proliferation. However, the investigated cationic liposomes showed more efficient delivery of siRNA, release of siRNA from endosomes and consequently had a more pronounced therapeutic effect on *in vitro* model of CML.

## 2. Materials and methods

### 2.1. Cell line

K-562 cells (obtained from the Institute of Cytology RAS, St Petersburg, Russia) are CML in myeloid blast crises. Cells were cultured in RPMI 1640 medium (Biolot, Russian Federation), supplemented with 10% fetal bovine serum and maintained at 37°C in a humidified incubator with 5% CO_2_.

### 2.2. siRNA

The sequence of siRNA, directed against the BCR-ABL transcript, has previously been published [25]. The sense and antisense sequences of siRNAs are as follows: 5’-GCAGAGUUCAAAAGCCCUUdTdT and 5′-AAGGGCUUUUGAACUCUGCdTdT, respectively. For the negative control we used a nonsense scrambled siRNA with the following sense and antisense sequences: 5′-CCGGUGUGCUUCGACAACUdTdT and 5′-AGUUGUCGAAGCACACCGGdTdT respectively. To evaluate the transfection efficiency, we used Cy3, FAM, ROX and JOE fluorescently labeled siRNA (5′-terminus of antisense strand was labeled). siRNAs were chemically synthesized (DNA-synthesis, Russian Federation).

### 2.3. Peptides

The sequences of the cell penetrating peptides EB1 (LIRLWSHLIHIWFQNRRLKWKKK-NH_2_) [15] and MPG-ΔNLS (ac-GALFLGFLGAAGSTMGAWSQPKSKRKV-cya) [26] were previously published. The peptides were synthesized by LLC “Verta” (Russia) with optical purity >97% (HPLC).

### 2.4. Cationic liposomes

A solution of 1,26-bis(cholest-5-en-3β-yloxycarbonylamino)-7,11,16,20-tetraazahexacosan tetrahydrochloride (2X3) [27] in a mixture of CHCl_3_–CH_3_OH (1:1 vol.) was added to a solution of 1,2-dioleoyl-*sn*-glycero-3-phosphoethanolamine (DOPE, Avanti Polar Lipids, USA) in CHCl_3_ at a molar ratio of 1:3 and gently stirred. A solution of DSPE-PEG_2000_ (Avanti Polar Lipids, USA) in CHCl_3_ was added to the lipid mixture and organic solvents were removed in vacuo, the lipid film obtained was dried for 4 h at 0.1 Torr to remove residual organic solvent and was hydrated for 60 min using deionized water (MilliQ, USA) at 24°C. The liposomal dispersion was sonicated for 15 min at 70–75°C in a bath-type sonicator (Bandelin Sonorex Digitec DT 52H, Germany), filtered (0.45 µm Chromafil_®_ CA-45/25; Macherey–Nagel, Germany), flushed with argon and stored at 4°C.

### 2.5. Complexes siRNA/peptide preparation

To prepare the complexes between EB1 and siRNA, we used various ratios of at molar excess at various ratios of 5:1, 10:1, 20:1, 30:1, and 40:1 (EB1: siRNA molar ratio), which correspond to charge ratio (CR) values of 1/1, 2/1, 4/1, 6/1 and 8/1, respectively. The solution of siRNA (20 pMol) in 2.5 µL nuclease-free water (Sigma, USA) was mixed with 2.5 µL of peptide’s solutions in nuclease-free water. The mixture was vortexed for 20 seconds, incubated for 20 minutes at 25°C, and then diluted with the required volume of water.

For transfection experiments, we formed complexes of EB1 with siRNA in nuclease-free water by mixing a siRNA solution (100 pMol or 20 pMol) and EB1 solution in equal volumes at a CR of 4/1, followed by vortexing for 10 seconds and incubation for 20 minutes at 25 ^0^C.

### 2.6. Complexes siRNA/cationic liposomes preparation

To measure the physicochemical characteristics of the *siRNA-cationic liposomes complexes* (lipoplexes) 20 pMol of siRNA in 2.5 µL nuclease-free water was mixed with 2.5 µL of cationic liposomes in nuclease-free water at CR of 1/1, 2/1, 4/1, 6/1, 8/1, 10/1. Then, complexes were incubated for 20 min at 25 ^0^C and diluted with the required volume of water.

For transfection experiments, complexes of cationic liposomes with siRNA were formed in RPMI medium (Gibco, USA) without serum by mixing of a siRNA solution (100 pMol or 20 pMol) and 1mM cationic liposomes solution in equal volumes at CR of 8/1 with vortex mixing for 10 s followed by incubation for 20 min at 25°C.

### 2.7. Size and zeta potential measurement

Solutions of the complexes were prepared as described in nuclease-free water. Particle size was determined by dynamic light scattering, and zeta-potentials were measured by electrophoretic method with the Zetasizer 3000 HS instrument (Malvern Instruments, UK). Results are expressed as the average of three measurements.

### 2.8. Atomic Force Microscopy (AFM)

Solutions of EB1 or liposomes were prepared as described above in nuclease-free water. To evaluate the sample surface topography, we added 30 µL of water to 5 µL of a peptide or liposome solution, or to 5 µL peptide/liposome complexes with siRNA and applied to a freshly cleaved mica (SPI Supplies, West Chester, PA, USA). The mixture was then incubated at room temperature for 1 minute. Afterwards, the solution was discarded, and the mica surface was washed with water and dried with a Concentrator Plus (Eppendorf, Hamburg, Germany). To measure sample surface topography, we used a SolverNext scanning probe microscope (NT-MDT, Zelenograd, Russia) in semicontact mode with an NSG03 probe (NT-MDT, Zelenograd, Russia). Images were processed using Gwyddion 2.62 software (Czech Metrology Institute, Brno, Czech Republic) [28].

### 2.9. Transfection

The day before transfection, cells were plated and incubate under normal growth conditions. On the day of transfection, cells were plated at a cell density 50 × 10^3^ cells in 24-well plate or 10 ^4^ cells in 100 µL in 96-well plate in complete RPMI 1640 medium. After a short period of cell incubation, we added 100 µL or 20 µL of EB1/bcr-abl siRNA complexes and 50 µL or 10 µL of 2X3-DOPE-PEG /bcr-abl siRNA complexes to the cell in 24- or 96-well plate (100 pMol or 20 pMol siRNA per well), respectively, and incubated the cells in a 37 ^0^C incubator until analysis.

### 2.10. Inhibition of endocytosis

In endocytosis inhibition experiments K-562 cells were incubated in serum-free medium for 2 hours before transfection. After that, cells were treated with endocytosis inhibitors: 100 µmol dynasore, or 10 µg/ml cytochalasin D, or 5 µg/ml filipin (Sigma, USA) [22], for 30 minutes at 37°C in serum-free medium. Cells were then incubated with the transfection complexes in the presence of the endocytosis inhibitors. After 15 minutes of transfection, cells were washed with PBS and analyzed by flow cytometry.

For nonspecific inhibition of endocytosis, K-562 cells were pre-maintained at 4°C for 30 minutes and then transfected with EB1/siRNA complexes for 15 minutes at low temperature (4°C) to block the cellular uptake via the endosomal pathways. Then the cells were then washed three times with cold PBS. The efficiency of transfection was then assessed by flow cytometry.

### 2.11. Flow cytometry analysis

The transfection of K-562 cells was carried out according to the earlier described method (sec 2.9). After the incubation time, the cells were centrifuged at 1500 rpm for 5 minutes and washed twice with PBS to remove any membrane-bound complexes. The supernatant was discarded, and the cell pellet was resuspended in 100 µL PBS. Fluorescence analysis was immediately performed using a CytoFLEX flow cytometer (Beckman Coulter, Indianapolis, IN, USA). Transfection efficiency was quantified using flow cytometry based on two parameters: percent of transfected cells (%) and mean fluorescence intensity of cell population (MFI). Percent of transfected cells is represented as the percentage of fluor-positive cells (JOE or ROX). The MFI was calculated as a median for fluor-positive cells. The results were analyzed in Kaluza software and were expressed as the mean and standard deviation obtained from three samples.

### 2.12. Laser confocal scanning microscopy

To analyze the uptake and localizations of fluorescent-labeled siRNA in cells, we used a confocal laser scanning microscope Leica TCS SP8 (Leica Microsystems, Germany) with an oil immersion objective 60X and 1.25 NA. The base plane of the cell was imaged with the scanning frequency of 400 and image size of 1024 × 1024 pixels.

For analysis, K-562 cells were transfected with JOE-labeled siRNA complexes as previously described. At 24 hours post-transfection, the cells were washed in PBS to eliminate free fluorescent siRNA molecules outside the cells, and then placed on poly-L-lysine-coated coverslips. The cells were immediately fixed in 4% paraformaldehyde and permeabilized in 0/1% Triton. Nuclei and F-actin were stained with DAPI and Alexa Fluor 680 phalloidin, respectively.

For transfection of K-562 cells with the EB1/siRNA complex labeled with ROX, we followed the same procedure as described earlier in the presence of LysoTracker ™ Blue DND-22 fluorescent acidotropic probes for labeling and tracking acidic organelles in live cells (L7525, ThermoFisher, USA). The staining was carried out according to the manufacturer’s instructions. We evaluate the results using confocal laser scanning microscopy. The images were processed and analyzed using ImageJ software (National Institutes of Health, Bethesda, MY, USA).

### 2.13. Immunofluorescence

The cells were transfected with complexes as described above, and then transferred to PBS at the indicated time-point. They were then placed on poly-L-lysine-coated coverslips and immediately fixed in 4% paraformaldehyde for 15 minutes at 24°C. After fixation, the cells were washed with PBS, permeabilized with 0.1% Triton X-100 for 15 minutes at 24°C, washed and blocked in 1% BSA for 30 minutes at 24°C to prevent non-specific antibody binding. Subsequently, the cells were incubated with the primary antibodies, consisting of Anti-EEA1 antibody - Early Endosome Marker (ab109110 Abcam, UK), Anti-RAB7 antibody (ab137029 Abcam, UK) and Anti-LAMP1 antibody - Lysosome Marker (ab24170 Abcam, UK) for one hour at 24°C after diluting them in 1% BSA. Primary antibodies were used at a dilution 1:1000. Following incubation, cells were washed with PBS containing 0.1% Tween-20 and then incubated with secondary antibodies GAM-Alexa Fluor 568 (Molecular Probes, USA) and DAR-Alexa Fluor 633 (Molecular Probes, USA) at a dilution 1:500 for one hour at RT. Finally, the cells were mounted in Fluoroshield Mounting Medium with DAPI (Sigma, USA).

The images were analyzed using confocal laser scanning microscope Leica TCS SP8 (Leica Microsystems, Germany) and processed and analyzed through the ImageJ software (National Institutes of Health, Bethesda, MY, USA). The quantitative co-localization analysis was performed using ImageJ JACoP Plugin to determine Manders’ co-localization coefficient M2[29]. This coefficient is defined as the sum of the intensities of the selected green objects containing red signal, divided by the sum of the intensities of all selected green objects. Results were represented as mean ± standard deviation.

### 2.14. RNA purification and Quantitative RT-PCR

mRNA levels of *BCR-ABL* gene after transfection were quantified by real-time PCR (RT-qPCR). RNA was isolated using the TRIzol™ Reagent (Thermo Scientific, USA) according to the manufacturer’s instructions. The extracted RNA concentration was then assessed via spectrophotometry (NanoDrop 2000c Spectrophotometer, Thermo Scientific, USA). The samples were pretreated with RQ1 RNase-Free DNase (Promega, USA). The reverse-transcription reaction was performed by RNAscribe RT (Biolabmix, Russia) following the manufacturer’s instructions. The RT-qPCR was performed employing SsoAdvanced universal SYBR Green supermix (BioRad, USA) for *BCR-ABL* and glyceraldehyde-3-phosphate dehydrogenase (*GAPDH*) with the following probes and primers: bcr-abl FP, 5′-TCCGCTGACCATCAATAAGGA-3′; bcr-abl RP, 5′-CACTCAGACCCTGAGGCTCAA-3′ [30]; gapd FP, 5′-CAGTCAGCCGCATCTTCTTTTGGCGTCG-3′; gapd RP, 5′-CAGAGTTAAAAGCAGCCCTGGTGACCAGG-3′. Using an CFX96 Touch Real-Time PCR Detection System (BioRad, USA), conditions for reaction mixture involved 45 cycles of 95 ^0^C denaturation and 60°C annealing. The analysis to determine differences in gene expression was performed using 2^−ΔΔCT^ method [31]. *BCR-ABL* CT was normalized against *GAPDH* CT, and the results are expressed as a relative quantity of the targeted mRNA.

### 2.15. Cell proliferation assay

The K-562 cells (10^4^ cells/well) were seeded in 96-well cell culture plates and then treated with the investigated complexes. Cell viability was assessed using the MTS [3-(4,5-dimethylthiazol-2-yl)-5-(3-carboxymethoxyphenyl)-2-(4-sulfophenyl)-2H-tetrazolium, inner salt] assay (Promega, UK). After the incubation periods, 20 µL of MTS was added to each well. The plates were then placed in the dark and incubated for 2 hours at 37°C. Subsequently, absorbance values (A) were obtained at 490 nm using a microplate reader (ClarioStar, BMG Labtech, Germany). The percentage of cell viability of the test sample was calculated by using formula: 100 × *A*_*test*_*/ A*_*control*_.

## 3. Results and discussion

### 3.1. Physicochemical characteristics of complexes

During the initial stage of this study, it was necessary to prepare and characterize the complexes of either EB1 or liposomes with siRNA to deliver into K562 cells. It is well established that size and surface charge of nanoparticles strongly influence their intracellular delivery efficiency [32][33]. Specifically, the size of these nanocomplexes must be less than 200 nm to facilitate cellular uptake and tissue distribution. Additionally, the net charge of such complexes is a critical parameter that influences siRNA binding and complex stability. The charge should be positive and sufficient to effective interaction between the complexes and the cell membrane, thus promoting internalization. However, the charge should not be too high, as this could lead to cellular toxicity and aggregation of the complexes with negatively charged serum proteins (such as albumins) [34][35]

To characterize the balance between peptide positively charged amino acid residues or cationic lipids amino groups and negatively charged RNA molecules, two similar parameters are commonly used for peptides and liposomes. The charge ratio (CR) is defined as the number of positive charges from the peptide or nitrogen atoms in the cationic lipid per negative charge phosphate groups from the siRNA. CR ratio is a critical parameter that can significantly influence the peptide or liposome/siRNA complex formation and efficiency of siRNA delivery.

The cell-penetrating peptide EB1 is a derivative of penetratin peptide. It has a total net charge +8 in physiological pH. EB1 is secondary amphipathic, which means it exhibits amphipathic properties upon acquiring an alpha-helical secondary structure on a membrane. Previously, we have shown that CR of complex formation is more critical than molar ratio and have identified optimal conditions for forming complexes between cell penetrating peptide and siRNA [36].To prepare the complexes between EB1 and siRNA, the peptide was added in a molar excess at various CR of 1/1, 2/1, 4/1, 6/1 and 8/1. The hydrodynamic size and zeta potential of these EB1/siRNA complexes at different CR were characterized using dynamic light scattering (DLS). The complexes, formed at CR of 4/1, 6/1 and 8/1, were found to have an optimal size of approximately 100 nm and a high net positive charge, which could facilitate their cellular uptake. A notable difference in zeta potential was observed among the complexes formed at CR of 1/1, 2/1 and 4/1. Additionally, the complex formed at 2/1 CR demonstrated significant differences in size compared to those formed at a 4/1 CR, indicating that this complex possessed high aggregation ability in water and was less effective for intracellular delivery than nanometer-sized complexes (Table1). Based on these findings, it can be concluded that the complexes formed at CR of 4/1 and higher exhibited optimal zeta potential and hydrodynamic size for efficient transfection *in vitro*. Thus, CR of 4/1 was established as an optimal ratio, and was used for the further experiments.

For our study, we prepared cationic liposomes based on polyacationic lipid 2X3 and helper-lipid DOPE with an increase in molar ratio towards the helper lipid DOPE. Specifically, we utilized 2X3-DOPE at a 1:3 molar ratio and modified liposomes with DSPE-PEG_2000_ (0.62% mol.). The encapsulation of siRNA primarily based on electrostatic interactions between anionic nucleic acids and 2X3. DOPE, a zwitterionic lipid, is added for its fusiogenic property, which promote endosomal escape of siRNA into the intracellular compartment [37]. Our previous findings demonstrate that an increase in the proportion of DOPE in liposome formulations, based on 2X3, enhances the efficiency of mRNA delivery to eukaryotic cells [38]. The addition of 1,2-distearoyl-sn-glycero-3-phosphoethanolamine-N-[methoxy(polyethyleneglycol)-2000] (DSPE-PEG_2000_) plays a significant role in achieving efficient nuclease protection of siRNA and prolonged drug release of the PEGylated lipoplexes [39]. The prepared cationic liposomes had a size of 59 ± 1 nm and ζ-potential of +50.9±1.3 mV, as characterized by DLS. The solution of cationic liposomes in water was stable, possessing the PDI value of 0.23 ± 0.01.

We prepared lipoplexes using CR ratios 1/1, 2/1, 4/1, 6/1, 8/1 and 10/1, and characterize their physicochemical properties. Based on our findings, lipoplexes created with an CR ratio of 8/1 demonstrated the most compact complex formation and exhibited high ζ-potential (as shown in Table 1).

**Table 1.** Characterization of EB1/siRNA or 2X3-DOPE-PEG/siRNA complexes based on size, polydispersity index (PDI), and zeta potential (ZP) values. Each value represents the mean ± standard deviation of three measurements.

| Charge Ratio (CR) | EB1/siRNA |  |  | 2X3-DOPE-PEG/siRNA |  |  |
| --- | --- | --- | --- | --- | --- | --- |
|  | Size (nm) | PDI | ZP (mV) | Size (nm) | PDI | ZP (mV) |
| 1/1 | 85 ± 1 | 0.24 ± 0.01 | - 41.4 ± 2.7 | 266 ± 9 | 0.39 ± 0.10 | - 15.0 ± 0.8 |
| 2/1 | 1226 ± 8 | 0.54 ± 0.03 | + 9.4 ± 1.6 | 209 ± 3 | 0.34 ± 0.01 | - 13.0 ± 1.1 |
| 4/1 | 105 ± 2 | 0.13 ± 0.01 | + 20.4 ± 2.5 | 243 ± 8 | 0.39 ± 0.01 | + 21.0 ± 3.5 |
| 6/1 | 85 ± 1 | 0.15 ± 0.01 | + 22.1 ± 3.0 | 195 ± 6 | 0.36 ± 0.02 | + 48.4 ± 1.0 |
| 8/1 | 78 ± 1 | 0.11 ± 0.01 | + 24.9 ± 2.0 | 168 ± 1 | 0.32 ± 0.02 | + 47.2 ± 1.9 |
| 10/1 | - | - | - | 218 ± 7 | 0.40 ± 0.01 | + 51.5 ± 0.40 |

To visualize the morphological shapes of complexes atomic force microscopy was performed. Based on the DLS results, we chose a CR of 4/1 for EB1/siRNA complexes and CR of 8/1 for 2X3-DOPE-PEG complexes. Figure 1 shows representative images of the surface topography of EB1/siRNA and liposome/siRNA complexes.

**Fig.1.**
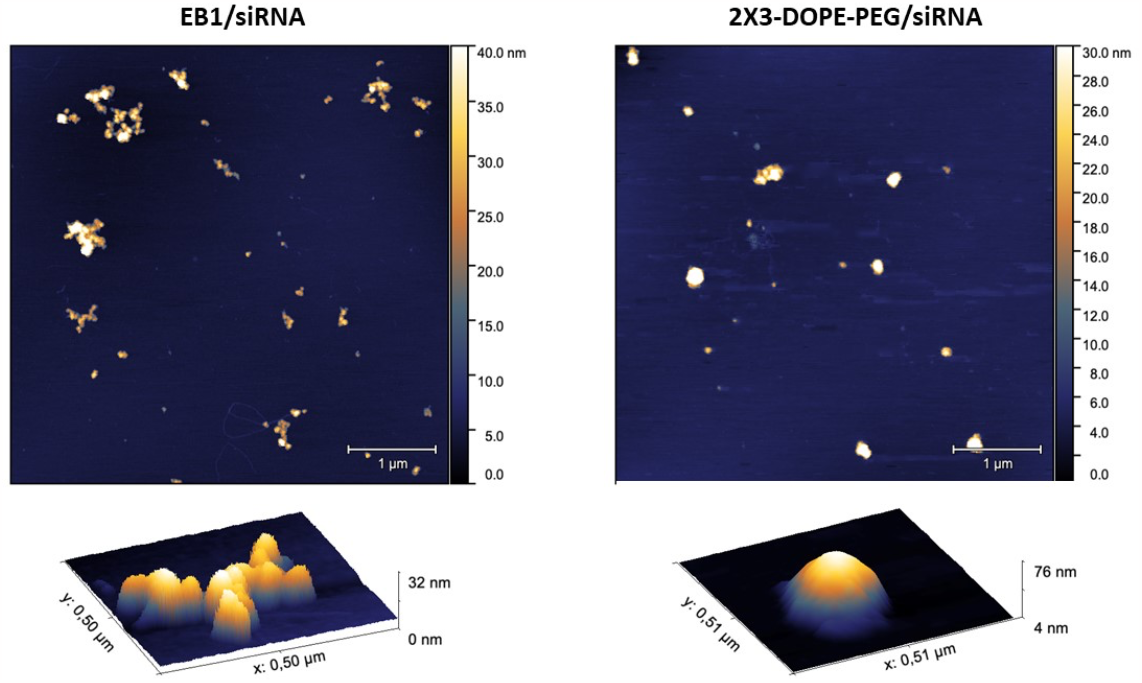
Surface topography of EB1/siRNA or 2X3-DOPE-PEG/siRNA complexes. The scale bar is 1 µm. Pseudo-color ruler on the right side of each image reflects the height distribution of particles in the sample, is located. The bottom row shows the characteristic particles of each sample in 3D view.

The 2D and 3D AFM images demonstrated that both EB1 and 2X3-DOPE-PEG were effective in condensing siRNA and forming particle-shaped nanocomplexes (see Figure 1). Additionally, the average size of 2X3-DOPE-PEG/siRNA complexes was found to be larger compared to the EB1/siRNA complexes.

Based on the analysis of physicochemical properties, we determined that the optimal CR of the peptide and siRNA was 4/1, while the optimal CR for cationic liposomes and siRNA was 8/1. These ratios were then utilized for *in vitro* experiments. Additionally, it was observed that the complexes formed between the peptide and siRNA have a smaller size and charges compared to lipoplexes.

### 3.2. Cytotoxicity study of EB1 and 2X3-DOPE-PEG in K-562 cells

To investigate the cytotoxicity of studied carriers for siRNA delivery we used concentrations of peptide and liposomes equivalent to complex formation with siRNA for transfection experiments. MTS assay was performed at 24, 48, 72, and 96 hours after adding the peptide or liposomes to K-562 cells. Our results showed that EB1 had a weak toxicity effect, whereas liposome addition led to a tendency of temporary metabolic activation of the cells at 24 hours and did not demonstrate any cytotoxicity (Figure 2).

**Fig.2.**
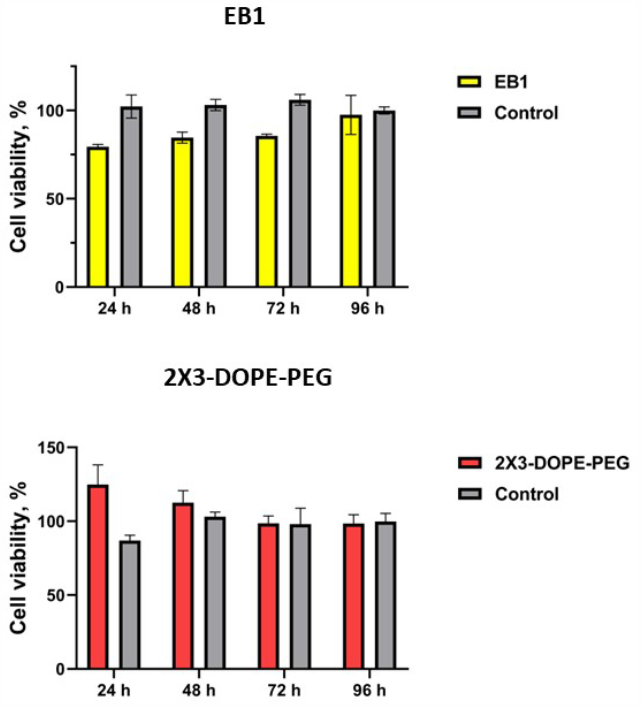
The cell viability of K-562 cells at 24, 48, 72, and 96 hours after treatment with EB1 and 2X3-DOPE-PEG. The mean and standard deviation of the measurements are presented based on three independent experiments.

The obtained data indicates that both investigated carriers, EB1 and 2X3-DOPE-PEG, are non-toxic and have the potential to be utilized as effective delivery vehicles for siRNA.

### 3.3. Cellular uptake efficiency of EB1/siRNA complexes in K-562 cells

The characterized complexes were evaluated for their ability to efficiently deliver fluorescently labeled siRNA into the K-562 cells (human chronic myeloid leukemia cell line). To evaluate the effectiveness of siRNA delivery mediated by liposomes or peptide, a flow cytometry assay was performed. After transfection with EB1/siRNA complexes at 4/1 CR and the 2X3-DOPE-PEG at 8/1 CR, almost equal percentage of cells internalized JOE-labeled siRNA after 24 hours of incubation were observed (83.6 ± 1.5% and 93.4 ± 0.4%, respectively). However, the mean fluorescence intensity (MFI) for EB1 (55266 ± 1750) was half that of 2X3-DOPE-PEG (115994 ± 5150) (Fig.3 a).

**Fig.3.**
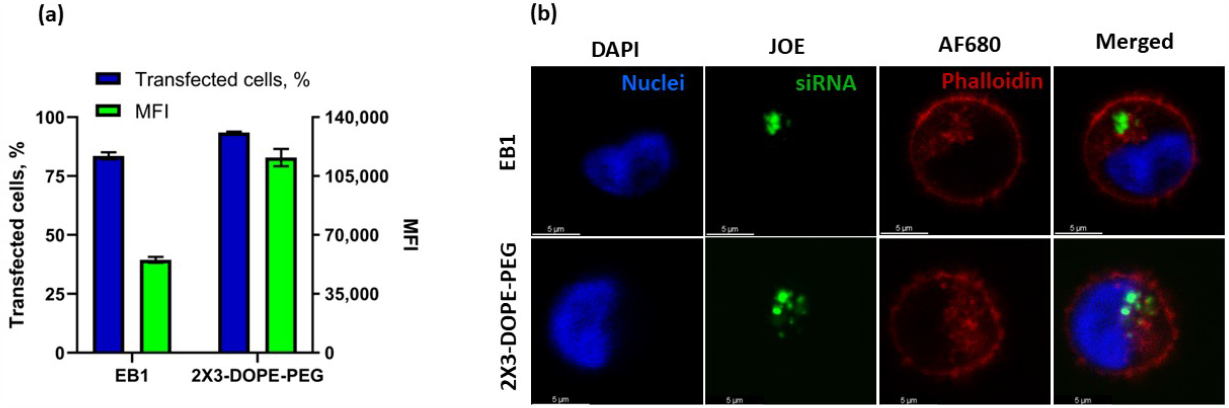
**(a)**Transfection efficiency of complexes in K562 cell. The flow cytometry analysis was performed 24 hours after transfection with JOE-labeled siRNA. The percentage of transfected cells is displayed on the left Y-axis, while the right Y-axis represents the mean fluorescent intensity (MFI). **(b)** Cellular uptake of complexes in K562 cells. Cells were transfected with siRNA using EB1 and 2X3-DOPE-PEG for 24 hours, and then analyzed by confocal microscopy. The image shows nuclear chromatin staining with DAPI (blue), JOE-labeled siRNA(green), and cytoskeleton staining with Alexa Fluor 680 phalloidin. The scale bar is 5 µm.

Confocal microscopy was used to study the subcellular localization of the complexes. According to confocal microscopy results, JOE-labeled siRNAs were internalized by the cells after 24 hours of transfection (as shown on Figure 3 b). The granular signals observed confirm the intracellular localization of studied complexes.

Both the flow cytometry and confocal microscopy data indicated that the investigated complexes efficiently mediated siRNA delivery into K-562 cells. 2X3-DOPE-PEG/siRNA complexes were found to deliver siRNA more efficiently compared to EB1/siRNA complexes.

### 3.4. Internalization mechanisms of the complexes in K-562 cells

Generalizing the current findings about mechanism of nanocomplexes’ cellular uptake remains challenging due to variation on the endocytic pathways that are dependent on cell types, delivered molecules, and different nanocomplexes [40]. Investigating the mechanisms of siRNA cellular uptake is crucial in determining intracellular transport and endosomal release, which are essential for safe and efficient therapeutic applications. Understanding the potential of delivery methods is also important in this regard.

Despite active research mechanisms by which CPPs are internalized into cells remain unclear. It is known that CPP can enter cells via direct translocation (energy-independent) or by endocytosis (energy-dependent). Studies have shown that CPPs and their cargoes can be taken up by cells through single or multiple endocytic pathways. The specific mechanism of siRNA delivery may depend on different experimental conditions such as CPP concentrations, cargo molecule characteristics, peptide-cargo complex properties, and the tissue or cell line type [41]. It has been proposed, that EB1 enters the cells via endocytosis but this has not yet been confirmed experimentally [15].

The main cellular process involved in the internalization of nanocomplexes is endocytosis, which includes macropinocytosis, clathrin-mediated endocytosis, caveolin-mediated endocytosis and clathrin/caveolae-independent endocytosis. To determine the endocytic pathway responsible for the uptake of the complexes, we have transfected cells with our complexes in the presence of different endocytic inhibitors: dynasore, an inhibitor of dynamin GTPase activity and blocks dynamin-dependent endocytosis in cells, cytochalasine D, an inhibitor of macropinocytosis and fillipin, known to inhibit caveola/raft-mediated endocytosis [42].

The transfection efficiency of siRNA by EB1 is decreased in the presence of endocytotic inhibitors, but not dramatically. It can be assumed that entry of EB1 into studied cells involves multiple cell entry pathways, including dynamin-dependent endocytosis, macropinocytosis and caveola-mediated endocytosis.

It was observed that treatment with fillipin did not affect the uptake when siRNA was transfected using 2X3-DOPE-PEG. However, treatment with dynasore and cytochalasine D resulted in a reduction of the uptake of the complexes (as shown on Figure 4). These findings confirm that the lipoplexes with siRNA primarily enter K-562 mostly through dynamin-dependent endocytosis and partially through macropinocytosis.

**Fig.4.**
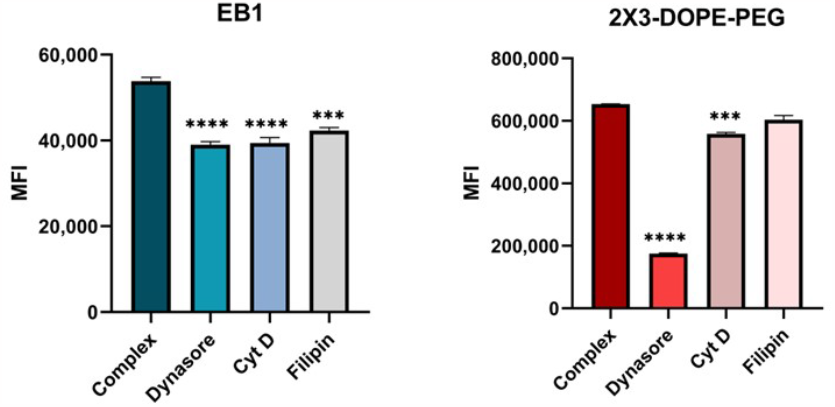
Mean fluorescence intensity of JOE (indicating siRNA uptake) measured by flow cytometry in K-562 cells. Cells were treated with JOE-labeled siRNA complexed with EB1 or 2X3-DOPE-PEG for 15 minutes in the presence or absence (‘Complex’) of endocytosis inhibitors including dynasore, cytochalasin D (‘Cyt D’) and filipin. The mean and standard deviation of the measurements are presented. The statistical analysis was performed using two-way ANOVA: ***—p < 0.001; **—p < 0.01; *—p < 0.05.

We additionally performed the transfection of K-562 cells by EB1/siRNA at 4 ^0^C to clarify the entry mechanism since almost all endocytic pathways are energy-dependent processes that can be inhibited at low temperatures and some CPP could enter cells via direct translocation (energy-independent). As a positive control of CPP that enters cells via an energy-independent membrane translocation mechanism, we used the well described MPG-ΔNLS peptide [26]. Previously we have demonstrated the optimal CR for stable MPG-ΔNLS/siRNA complex formation and delivery efficiency [36]. Our results showed that transfection efficacy of siRNA by EB1 at 4 ^0^C for 15 minutes was significantly reduced (7.1 ± 0.1%) compared to cells incubated at 37 °C (80.2 ± 0.8%) as measured by flow cytometry (Figure 5), while the efficiency of transfection with the MPG peptide did not decrease significantly. This indicates that the internalization of investigated EB1/siRNA complexes into K-562 cells most likely occurred via energy-dependent processes.

**Fig.5.**
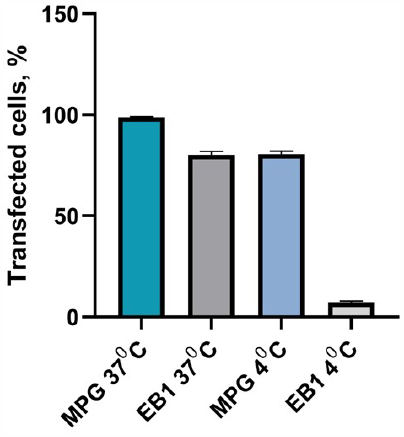
Transfection efficiency of EB1/siRNA-ROX complexes in K-562 cells at low temperature compared with that of the MPG/siRNA-ROX complexes. The bar graphs show the percentage of ROX-positive live cells 15 minutes post-transfection.

We also investigated the transfection of K-562 cells after incubation with EB1/siRNA-ROX complexes for 15 minutes in the presence of LysoTracker ™ Blue DND-22. We examined the colocalization of ROX-labeled siRNA with acidic cell compartments stained with fluorescent dye LysoTracker ™ Blue DND-22. Figure 6 demonstrates colocalization of delivered siRNA with fluorescent dye, that specifically stains acidic compartments within live cells. Thus, suggesting that EB1/siRNA complexes were internalized into the K-562 cells by endocytosis and localized in endocytic compartments with acidic pH. Previous research has shown that endosomes in K-562 cells are more acidic than those in other cell types [16].

**Fig.6.**
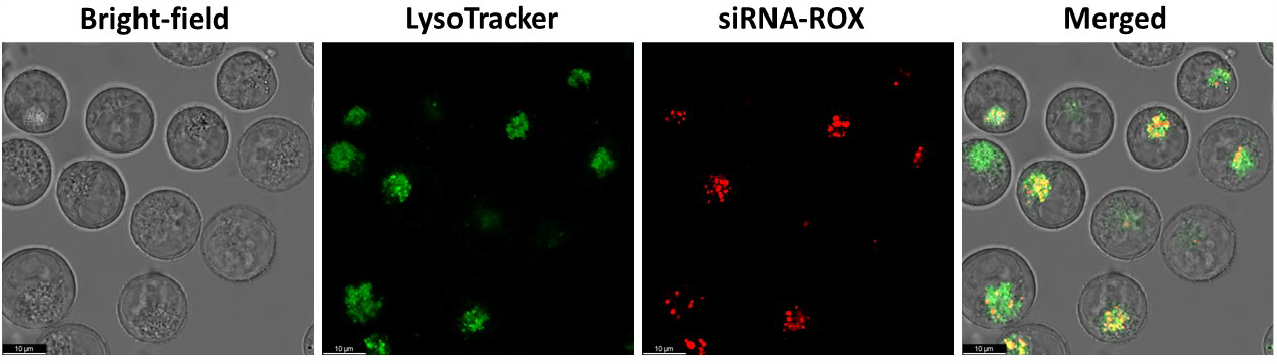
Live imaging confocal microscopy of the cellular uptake of peptide EB1/siRNA complexes in K-562 cells, with costaining using LysoTracker. LysoTracker ™ Blue DND-22 is stained green, while siRNA-ROX is in red. The scale bar is 10 µm.

These findings confirm that EB1/siRNA complexes enter cells through various mechanisms of endocytosis, including dynamin-dependent endocytosis, micropinocytosis, caveola-mediated endocytosis, and possibly dynamin-independent endocytosis.

### 3.5. Endosomal escape study

As we have demonstrated in the pervious step, the nanocomplexes are internalized into K-562 cells through the endocytosis pathway. Internalization via endocytosis involves entry into an endocytic vesicle, fusion with the early endosomal compartment (EEs), maturation into a late endosome (LEs) and accumulation in the lysosome (LYs). One of the limiting factors in achieving effective siRNA-based therapy is facilitating endosomal escape and provide cytosolic delivery of the therapeutic siRNA. Failure to escape the endosome leads to entrapment and possible degradation in the LYs.

CPPs have been widely used for the siRNA delivery but the association loci on cell surface, cell internalization pathways, intracellular trafficking, and escape from endosomes are still a topic of debate [43]. We have shown that the entry mechanism of EB1 and siRNA complexes into K-562 cells may involve several pathways of endocytosis. EB1 is an analog of penetratin derived from the Antennapedia protein homeodomain, with some amino acids replaced with histidines and N-terminally extended to potentially penetrate the endosomal membrane. EB1 demonstrated better RNAi-mediated gene silencing compared to its parent peptide penetratin in HeLa and HepG2 cells, probably due to the its improved ability to form complexes with siRNA and promote endosomal escape [15]. However, the mechanism of cell entry and release of delivered siRNA from endosomes has yet to be fully investigated.

Lipid nanoparticles (LNP) enter the cells via endocytosis with approximately 95% of LNPs being endocytosed within 30 minutes. However, only a small amount of the siRNA delivered by LNPs can escape the endosomes and reach the cytosol [44]. Cationic liposomes 2X3-DOPE-PEG contain the helper lipid DOPE, which facilitates the escape of nucleic acids from endosomes, and is simultaneously modified by PEG, that might overcome potential aggregation of cationic liposomes and enhance gene transfection efficiency *in vivo*. However, this represents a major barrier for cationic liposomes’ internalization and nucleic acid endosomal escape, resulting in significantly reduced transfection efficiency [45]. Therefore, studying the endosomal escape of delivered siRNA is a crucial step in assessing the prospects of delivery systems.

To investigate the poorly understood and inefficient process of endosomal release of delivered siRNA, we examined the uptake and endosomal escape of siRNAs formulated in nanocomplexes using confocal microscopy of immunolabeled K562 cells. Colocalization analysis was employed to determine if labelled proteins colocalize with the investigated labelled structures. We examined the colocalization of fluorescent-labeled siRNA with marker proteins for specific endomembrane compartments, including EEA1 (early endosomes, EEs, marker), Rab7 (late endosomes, LEs, marker) and LAMP1 (lysosomes, LYs, marker).

Our findings show that, following transfection of K-562 cells with EB1/siRNA complexes, siRNA was present inside EEA1-positive vesicles 15 minutes after being added to cells (58%). We then observed the progression of siRNA through EEs to LEs and LYs with almost the same level of colocalization with the corresponding markers (∼60%). This suggests that escape of siRNA from LEs in case of delivery by EB1 is limited (Figure 7).

**Fig.7.**
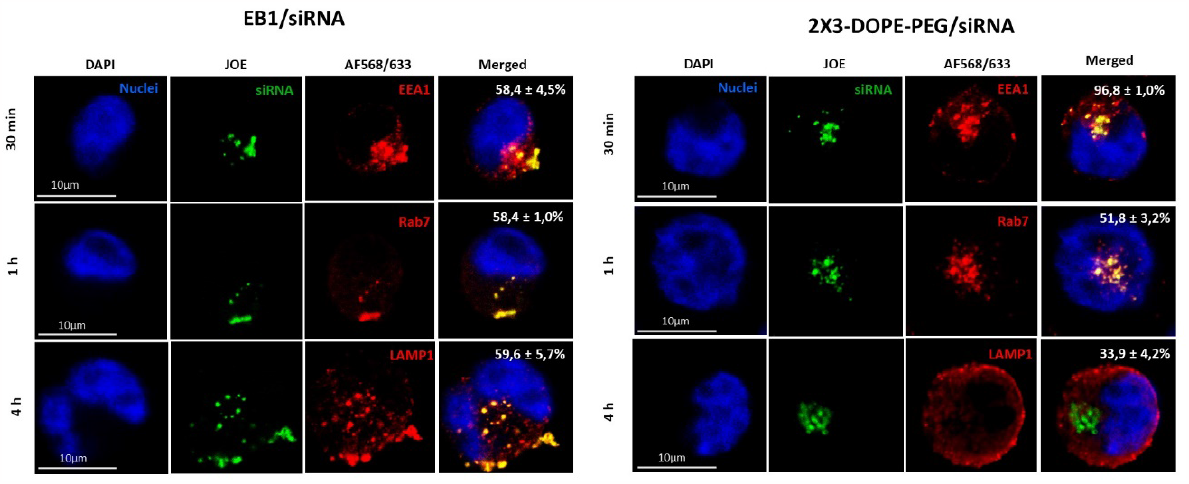
Confocal microscopy of immunolabeled K-562 cells transfected with EB1/siRNA or 2X3-DOPE-PEG/siRNA complexes using anti-EEA1, anti-Rab7 and anti-LAMP-1 antibodies (shown in red). Following transfection for 30 minutes, 1 hour and 4 hours of transfection, cells were fixed, permeabilized and immunostained. Endosomes were labeled with EEA1 (Alexa Fluor 568, red) and Rab7 (Alexa Flour 633, red) markers, which indicated the presence of siRNA in early and late endosomes, respectively, while the LAMP-1 marker (Alexa Fluor 568, red) showed the localization of siRNA in lysosomes. The siRNA itself was labeled with JOE (showed in green). Quantitation of colocalization (% Colocolization) was performed as described in the methods. The data correspond to mean ± SD, n = 30 in the pictures. The scale bar is 10 µm

Confocal images obtained after transfection with siRNA/liposomes complexes show that siRNA is initially presented in EEs, followed by progression to LEs and LYs. By examining the colocalization of siRNA with EEA1, Rab7 and LAMP-1 marker proteins, we determined that the maximal colocolization of siRNA with the EEs markers EEA1 occurred 15 minutes post-transfection (96,8%). After 1 hour post-transfection siRNAs are partially located in LEs (51,8 %) and after 4 hours post-transfection the colocalization of siRNA with LYs markers LAMP-1 was low (33,9 %) (Figure 7). These results indicate that siRNA passes through the EEs and LEs compartments and accumulates minimally in lysosomes, suggesting that siRNA may escape from LEs.

Our findings suggest that cationic liposomes enter cells primarily through dynamin-dependent endocytosis and efficiently release the delivered siRNA molecules. It should be noted that liposome modification with DSPE-PEG_2000_ (0.62% mol.) does not inhibit lipoplex internalization and siRNA endosomal escape. However, the entry mechanism of the peptide EB1 is more complex, and we observed that the delivered and visualized siRNA molecules are present at the same level in EEs, LEs and LYs. This may indicate limited siRNA release from endocytic vesicles or a rapid partial release from EEs that we did not detect.

### 3.6. The efficiency of target gene silencing (RT-PCR)

Our study has demonstrated that the nanocomplexes we investigated efficiently deliver siRNAs and release them from lysosomes into cells. The extent of this release is more pronounced in the case of cationic liposomes. Further, we evaluated the efficiency of silencing delivered siRNAs against the target gene *BCR-ABL*. The standard method for assessing the effectiveness of siRNA-based nanocomplexes is the quantitative measurement of gene silencing activities of siRNA. We measured *BCR-ABL* expression in K-562 cells relative to the housekeeping gene *GAPDH* by RT-PCR and calculated the quotient of *BCR-ABL/GAPDH*. Our results, as shown in Figure 8, demonstrate a significant decrease in BCR-ABL mRNA levels after transfection of K-562 cells with anti-bcr-abl siRNA mediated by peptide and liposomes. Transfection with EB1/siRNA complexes led to a maximum reduction of BCR-ABL mRNA at 24 hours (72.5 % after transfection), which gradually decreased over time. However, when using 2X3-DOPE-PEG/siRNA complexes, we observed a reduction of the BCR-ABL mRNA level by 90.5 % at 24 hours after transfection, with the effect persisting until 72 hours. Specifically, we found that after 48 hours and 72 hours, the reduction was 93.6% and 91.6%, respectively.

**Fig.8.**
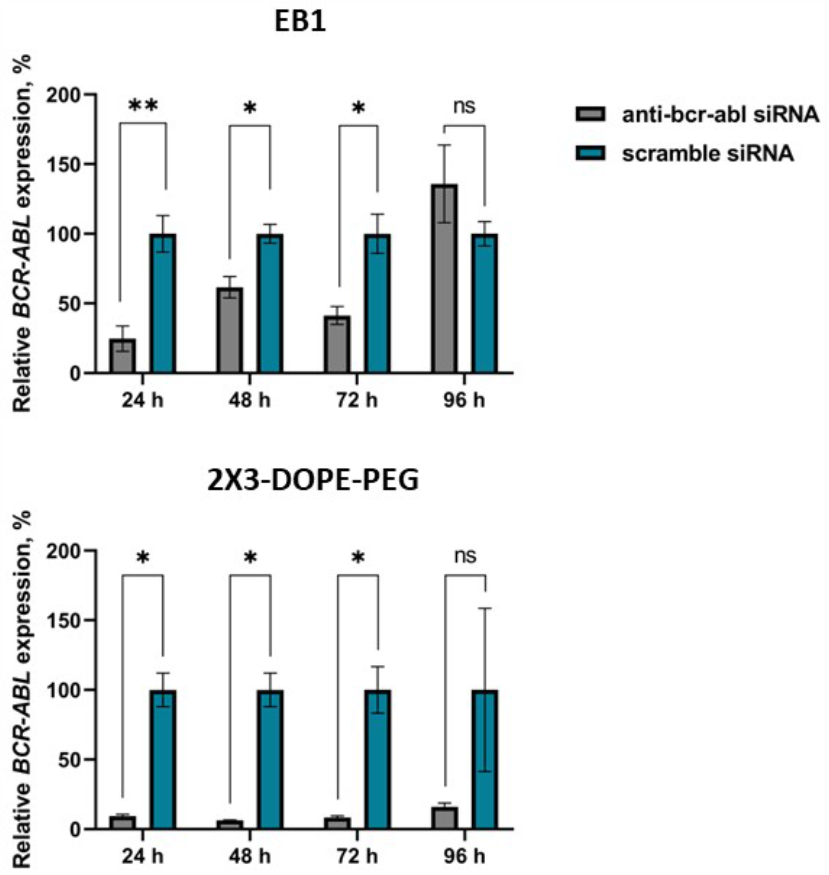
The results of the RT-PCR analysis showing BCR-ABL mRNA levels in K-562 cells at 24, 48, 72 and 96 hours after transfection with EB1/anti-bcr-abl siRNA complexes or 2X3-DOPE-PEG/anti-bcr-abl siRNA. The statistical analysis was performed using two-way ANOVA: **—p < 0.01; *—p < 0.05. p values > 0.05 (not significant – ‘ns’) are marked. To calculate fold change, the mean gene expression in samples treated with complexes containing nonsense/scrambled siRNA was considered as 100%.

Our results indicate that the transfection of K-562 cells with complexes of the peptide and cationic liposomes with siRNA leads to effective silencing of the *BCR-ABL* gene. However, using the EB1-based complexes leads to a less pronounced effect of inhibition of *BCR-ABL* gene expression. This could be due to a higher dose of siRNA entering the cytoplasm when delivered by cationic liposomes, which correlates with the data obtained in cellular uptake efficiency and endosomal escape studies. Furthermore, it could be suggested that 2X3-DOPE-PEG/siRNA complexes are more stable in the culture medium, which may enhance the delivery efficiency over time compared to peptide-based nanocomplexes due to modification by DSPE-PEG.

### 3.7. Growth Inhibition by BCR-ABL Silencing

The protein product of *BCR-ABL* gene is fusion oncoprotein which exhibits deregulated tyrosine kinase activity and is involved in the development of uncontrolled proliferation of CML cells. The effects of *BCR-ABL* silencing on abnormal proliferation of cells were evaluated by measuring cell viability after transfection using the MTS assay. It was shown that complexes of 2X3-DOPE-PEG with therapeutic siRNA significantly inhibited cell proliferation, with the highest levels at 72 hours (∼70%). In contrast, complexes of EB1 with the anti-bcr-abl siRNA demonstrated maximum reduction in proliferation after 24 hours post-treatment (∼50%), a gradual decrease in the antiproliferative effect over time (Figure 9). These findings are consistent with RT-PCR results.

**Fig.9.**
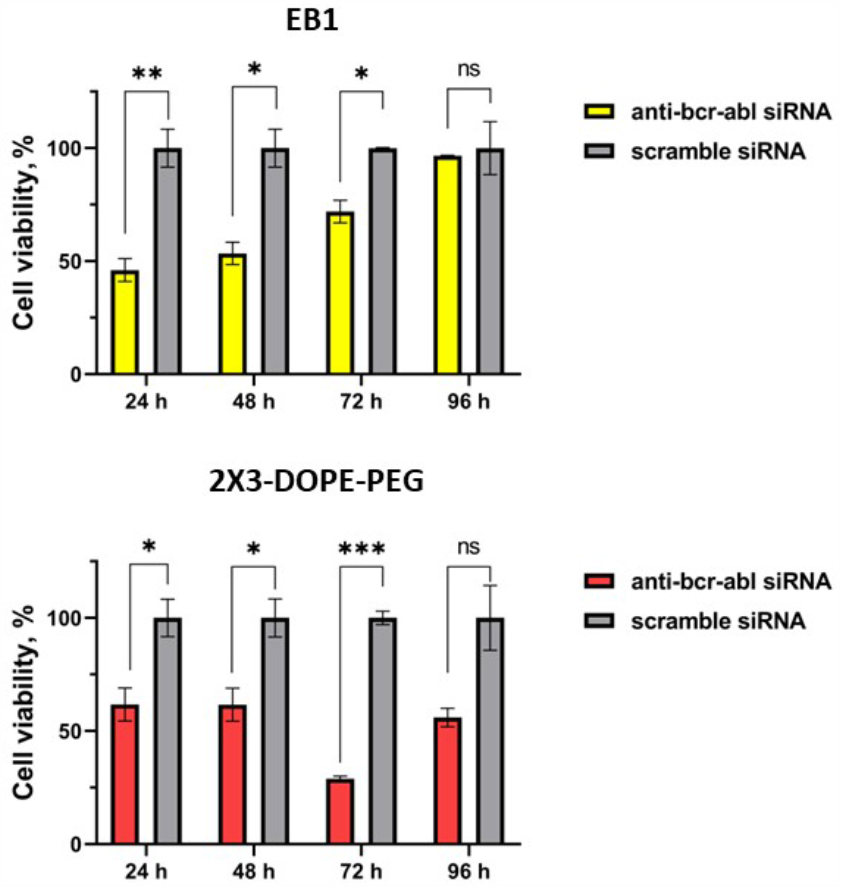
The viability of K562 cells after 24, 48, 72 and 96 hours of treatment with EB1/anti-bcr-abl siRNA and 2X3-DOPE-PEG/anti-bcr-abl siRNA treatment. The mean ± SD values are shown based on three independent experiments. The control is presented as transfection of siRNA with a nonsense/scrambled sequence. The statistical analysis was performed using two-way ANOVA: ***—p < 0.001; **—p < 0.01; *—p < 0.05. p values > 0.05 (not significant – ‘ns’) are marked.

The mechanism of internalization of the complexes and further intracellular transport of delivered siRNA molecules determine the therapeutic efficacy. The lower release of siRNA from endosomes leads to its degradation in LYs and insufficient silencing of the target gene [44]. Therefore, for EB1/siRNA complexes, we observe both a higher inhibition of mRNA level and a higher decreasing the number of living cell compare to liposome/siRNA complexes at 24 hours after transfection. In the case of liposome/siRNA, effective release into the cytosol is immediately reflected in mRNA expression level, which do not exceed 10% already by 24 hours, persist for 72 hours, and only slightly increases by 96 hours. According to cell viability assay results, we also observe a gradual decrease of cell proliferation by 72 hours, which is more pronounced compared to EB1/siRNA treatment. The subsequent increase in mRNA level and cell proliferation by 96 hours may be due to transient nature of the gene silencing effect and relatively long half-life of the BCR-ABL protein, the minimum level of which is sufficient to promote proliferation [46].

Our study has demonstrated that transfection of K-562 cells with the studied complexes leads not only to effective suppression of the target oncogene but also effectively reduces the pathological proliferation of tumor cells. Moreover, complexes with cationic liposomes were found to be more effective.

## 4. Conclusion

Chronic myeloid leukemia which is characterized by the presence of the *bcr-abl* fusion gene, provides an appropriate system for investigating gene therapy using various gene delivery vehicles. Several siRNA-based approaches have been developed to target the BCR-ABL fusion protein, including liposome-based delivery methods[47][48][49][50] and CPP [51]. However, the challenge of delivering siRNA for clinical implementation remains unresolved. Lipid nanoparticles (LNPs) have shown promise in efficient delivery of siRNA to the liver, as evidenced by the approval of Patisiran (ONPATTRO™) for the treatment of transthyretin-mediated amyloidosis based on LPN delivery. Several studies have reported siRNA encapsulated LNPs for treating various diseases such as cancer, viral infection, inflammatory neurological disorder, and genetic diseases. Nevertheless, delivering siRNA to other organs, notably hematopoietic tissues, remains a challenge. In the future, treatment with siRNA, alone or in combination with TKI could prove effective as therapeutic agents, reducing the risk of developing TKI-resistant mutants and potentially extending the drug’s efficacy period in CML patients [9].

In our study, we successfully compared two different nonviral delivery systems for anti-bcr-abl siRNA in CML K-562 cells. Our findings demonstrated that both approaches achieved efficient intracellular siRNA delivery, but cationic liposomes 2X3-DOPE-PEG exhibited a higher efficiency. We have shown earlier that the same liposomes 2X3-DOPE without DSPE-PEG_2000_ effectively deliver model mRNA *in vitro* [38]. In this work we demonstrated that PEG-modified liposomes provide high uptake efficiency, endosomal escape of siRNA that led to effective target gene silencing and antitumor effect, and PEG-modification, commonly used to improve *in vivo* efficiency does not impair efficiency *in vitro*. Therefore, liposomes 2X3-DOPE-PEG can be recommended as effective delivery vehicles for the future investigation of siRNA-based therapeutic strategy on *in vivo* CML model.

## Author Contributions

V.V., Y.Z.: Conceptualization, Investigation, Writing - Original Draft, Writing - Review & Editing, Visualization; A.E., V.K., N.K., I.T., A.B., O.D., M.E.: Investigation; M.M.: Conceptualization, Project administration, Writing - Review & Editing; A.V., M.D.: Conceptualization, Project administration, Funding acquisition.

## Acknowledgements

Darya Makarova for liposomes preparation.

